# Effects of generations in captivity and elevated rearing temperature on Ontario hatchery brook trout (*Salvelinus fontinalis*) fry quality and survival

**DOI:** 10.1101/2022.10.01.510397

**Authors:** A.S. Wilder, C.C. Wilson, T. R. Warriner, C.A.D. Semeniuk

## Abstract

With increasing environmental temperatures causing concern for the status of freshwater fishes, captive breeding programs may become increasingly important for conservation efforts as well as to support fisheries. Although captive broodstocks provide reliable gamete sources for production stocking, prolonged generations under hatchery conditions selection for hatchery conditions (domestication) and reduced phenotypic plasticity to novel environmental stressors. We assessed the effects of rearing temperature and number of generations spent in captivity on the survival and quality (indicated by lack of malformations) of long-term (F_20+_) and newly-captive (F_1_) strains of Ontario hatchery brook trout (*Salvelinus fontinalis*) with shared genetic history. We found that elevated temperatures decreased likelihood of survival between the hatched and fry stages. Additionally, we found that elevated temperature reduced fry quality of F_1_ fish whereas F_20+_ fish were less thermally sensitive, suggesting no reduction in plasticity due to captivity. The combined effects of elevated rearing temperatures and number of hatchery generations suggest that selection for captivity can occur rapidly (in one generation) even under benign conditions, and that additive stressor effects of captivity and temperature may impact newly established strains.

## Introduction

Climate change has been predicted to increase air and water temperatures globally (Creed et al., 2017), with freshwater aquatic ecosystems particularly vulnerable because of their dependency on air and precipitation temperatures combined with higher levels of human use (relative to size) compared to marine systems (Bunn, 2016; Cook, Burness, et al., 2018; Hanna et al., 2018; Potts et al., 2021; Stitt et al., 2014; Woodward et al., 2010). Effects are especially impactful on cold-water adapted freshwater species, including the economic- and culturally important salmonid group of fishes which have shown declines in abundance that are 64% higher than projected due to a combination of elevated water temperatures and loss of suitable habitat (Creed et al., 2017; Desforges et al., 2022; Lake et al., 2000; Myers et al., 2017). Specifically, elevated temperatures have been shown to impact the reproductive performance and survival of salmonids via multiple mechanisms: increased metabolic rates forcing energetic reallocation to homeostasis versus reproduction (Fenkes et al., 2016); the inhibition of production of hormones required for gonadogenesis reducing the overall reproductive potential of fishes (King et al., 2007; Pankhurst & King, 2010); and accelerated and subsequent underdeveloped hatch state of eggs (Baird et al., 2002; Hayes et al., 1953; McDermid et al., 2012). As embryonic and reproductive stages are the most thermally sensitive, increased temperatures pose a threat to reproduction and recruitment in cold-adapted salmonid populations (Dahlke et al., 2020).

Captive breeding of freshwater fishes has been employed as a common restoration tool to reverse or mitigate population declines (Cote et al., 2021; George et al., 2009; Houde, Garner, et al., 2015) particularly in freshwater fishes (Ward et al., 2008) such as salmonids (Solar, 2009).. Captive breeding programs require the cultivation of breeding adults through assigned mating, highly controlled environments, and strict nutrition to produce high quality offspring which are eventually used for stocking events (Bromage et al., 1992; Carrillo et al., 2000; Kuciński & Fopp-Bayat, 2021; Morissette et al., 2018; Olk et al., 2019). Progeny quality is quantified using egg fertilization, hatching success, survival through to the emergent fry stage (as defined by the period before the first exogenous feed), and fry quality as determined by the absence of malformations (Bromage et al., 1992; Carrillo et al., 2000).

Negative fitness consequences of captivity through domestication may be unavoidable despite extensive control measures throughout breeding and rearing (Milla et al., 2021). Because captive populations generally have greater survival (85-95%) than those in the wild (1-5%), recessive conditions may accumulate in turn may lead to greater instances of malformations as many developmental anomalies commonly seen in hatcheries are thought to have a genetic basis (Branson, 2008; Ihssen, 1978).

Despite these reproductive issues, hatchery broodstocks may inadvertently undergo multiple generations of selection to perform optimally in captive settings as a result of genetic, physiological, and behavioural adaptations to hatchery conditions (Liu et al., 2015). When compared to wild (F_0_) strains that have been acclimated in fish farms, individuals with greater number of generations in captivity can exhibit greater performance in fertilization success and egg survival (Liu et al., 2015).

Long-term adaptation to captivity can result in captive individuals losing plasticity in their responses to ‘irrelevant’ stressors (Mason et al., 2013; Milla et al., 2021). Reduced tolerance of hatchery salmonids to variable climatic temperatures has been documented, resulting in overall decreased survival of salmonid eggs when challenged with temperature stressors (e.g., *Oncorhynchus* spp., Murray and McPhail, 1988). To redress the losses in genetic and/or phenotypic plasticity of captive fish, individuals from the wild are introduced to begin a new broodstock strain, or crossbred with captive strains to infuse new genetic resources (Frankham, 2008). This action would serve to provide relief from domestication selection, reduce inbreeding potential, minimize effects of kinship, and help to prevent outbreeding depression upon stocking (Fisch et al., 2015). However, there is evidence that cold-freshwater fishes can experience selection in captive environments and exhibit reduced plasticity in as little as one generation (Christie et al., 2012, 2016; D. J. Fraser et al., 2019).

In addition to increasing genetic diversity to maximize post-stocking success (Rodewald, 2013), hatchery programs can also be accompanied by manipulating or enriching hatchery environments for broodstock and production fish. For addressing impacts of climate change on cold-water fish species, hatcheries provide opportunities for experimental evaluation of elevated rearing temperatures on offspring survival (McDermid et al., 2012; Molony et al., 2004; Pankhurst & King, 2010) and quality (Fjelldal et al., 2016; T. W. K. Fraser et al., 2015; T. S. Yamamoto et al., 1996). Furthermore, elevated temperatures have also been used to induce certain malformations (such as twinning) in instances where these specific developmental conditions are being explicitly studied (e.g., Atlantic salmon (*Salmo salar*) x Arctic charr (*Salvelinus alpinus*) hybrids; Fjelldal et al., 2016). Lastly, elevated rearing and incubation temperatures are also used to determine whether fish can be “phenotypically programmed” to deal with increased stressors upon release (e.g. climate change stressors, Cook et al., 2018b; Roberts et al., 2014). From a practical standpoint, elevated temperatures are also used by hatchery managers to increase egg survival and fry growth rates and therefore decrease rearing times (Baird et al., 2002; Marten, 1992), as well as manipulate hatch times (Weber et al., 2016) to increase production efficiency. Despite possible benefits, this thermal stress may cause a reduction in survival and increase in temperature-related malformations (e.g. blue sac disease in several charr species (Ihssen, 1978), spinal deformities in Atlantic salmon (T. W. K. Fraser et al., 2015), and possibly negate the advantages of priming and/or accelerating growth (e.g., carry-over effects at later-life history stages post stocking; Harbicht et al. 2020).

Most studies on egg survival and quality have focused on comparing responses of wild-type individuals with either short-term or long-term captive strains (Milla et al., 2021), By contrast, relatively few studies have compared newly-captive and long-term captive individuals; those that do tend to focus on post-release phenotypes rather than survival within a hatchery setting (Kostow, 2004; although see Howell et al. 2022). Since early development is an exceptionally vulnerable life history stage for fishes, particularly for temperature stressors (Dahlke et al., 2020), characterizing the integrated effects of generation times and performance under stress of current hatchery strains during early life can aid in predicting reproductive output and contributions to populations upon stocking (Haponski & Stepien, 2016). Here we used a two-by-two experimental design to investigate the effects of elevated rearing temperatures and number of generations in captivity on newly-captive (F_1_) and long-established (domesticated, F_20+_) hatchery strains of brook trout (*Salvelinus fontinalis*) with shared ancestry. Specifically, we compared how the domesticated and recent strains fared under normal hatchery conditions (i.e., captivity optimization) versus a stressful environment (i.e., loss of plasticity), assessing egg-survival and fry quality traits in both strains in benign and temperature-stressor environments. We predicted that elevated rearing temperatures would significantly reduce the likelihood of survival in both captive strains, with domesticated fish being more thermally sensitive. We also expect that while temperature will increase malformation rates, domestic fish will show higher instances of malformations and lower quality than the F_1_ strain under both temperature treatments due to unintentional inbreeding and suboptimal phenotype accumulation that is known to occur in hatchery settings (Crozier et al., 2011). Should domestication begin to act within one generation, we should expect to see no variation in survival between long-term captive and newly captive fish populations under normal captive conditions.

## Methods and Materials

### Fish origins

We used fish from two brook trout strains held at the Ontario Ministry of Natural Resources and Forestry (OMNRF) Codrington Fisheries Research Facility. The Hill’s Lake strain is a provincial production broodstock has been in the Ontario hatchery system for over 20 generations with some mid-20^th^ century genetic input from wild sources (OMNRF, 2005). The Scott Lake strain is a newly captive wild-origin research strain sourced from Scott Lake (5.48557, -78.7237) in Algonquin Park, Ontario. The wild brook trout population in Scott Lake was repeatedly stocked with the Hill’s Lake strain through the 20^th^ century (OMNRF unpubl. data) and shows extensive genetic introgression from the historical stocking (Harbicht et al., 2014) This strain had been in captivity for one full generation before the start of our experiment (i.e., parents were eggs that had been spawned from wild adults and brought into the hatchery system). Both strains are kept under common rearing conditions at the OMNRF Codrington Fisheries Research Facility (Codrington Ontario, 44.15°N, 77.80°W).

### Breeding and rearing design

Fertilized eggs were obtained from spawning adult fish at the Codrington Fisheries Research Facility. Between November 15 and November 29, 2019, ten male and 10 female four-year old adult Brook trout were used from the Hill’s Lake broodstock, >20 captive generations and considered domesticated. Ten male and 10 female adult trout of the same age were used from an F_1_ generation stock (parents of these fish were the last wild-reared generation) sourced from Scott Lake in Algonquin park. Adults were reared at ambient climatic conditions, using a flow-through ground water system, with an average water temperature of 7.17°C, current to wild habitat temperatures (see Howell et al., 2022). Five two-by-two fertilization crosses were performed within each population (i.e., five replicates of two females each crossed with two males). No siblings were used in the same 2×2 and no identical crosses were used. To collect gametes, adults were anesthetized using MS222 (Tricaine®-S Topical Anesthetics). Each adult had their pre-spawn mass recorded and photo taken, including a small tail-fin clip to allow us to determine parentage of offspring if so needed. Eggs were dry spawned into two 500mL glass jars by applying slight abdominal pressure to the female and fertilized at the hatchery. For the 2×2 cross, milt from two males was expressed via slight abdominal pressure and collected using disposable pipettes and partitioned across two females’ jars (i.e., there was no pooling across males or females). Hatchery water was used to activate eggs and milt; fertilization was stimulated by gently swirling the jar of eggs. Jars were then left to begin water hardening for approximately 10 minutes. Eggs were then disinfected with a 50mg/L solution Ovadine® according to OMNDMNRF protocol (Community Hatcheries Program, 2019), allowed to water harden for half an hour, rinsed with hatchery water three times, and transported back to the University of Windsor (Great Lakes Institute for Environmental Research aquatics facility) in coolers on ice. Upon arrival eggs were disinfected a second time, using dechlorinated Culligan® filtered water, following OMNDMNRF procedure before being placed into the incubation stacks. Families were equally split by equal volume into four egg cups, two replicates per temperature treatment (mean: 463 ml ± 79.456, Range: 341-627 ml; See Appendix A for sample sizes), that were randomized for position in the incubation stacks. The number of eggs transferred per family was not consistent across all females and families, as the female’s entire volume of eggs was used, and egg size was variable. Eggs were fertilized in three spawning batches across 3 weeks based on availability of spawning adults.

Within each strain, offspring were raised on two distinct temperature regimes: 7°C - near the optimal temperature for Brook trout eggs (Hokanson et al., 1973) and similar to temperatures at the Codrington hatchery (Howell et al., 2022), but within the physical capabilities of our rearing system, and 10°C - as this difference has been found previously to produce significant survival differences in Brook trout without approaching the TL50 of 12.7°C (Hokanson et al., 1973). An increase of 3°C also represents future water temperatures in accordance with current climate change projections (Austin & Colman, 2007; Trumpickas et al., 2009; Warriner et al., 2020). Each temperature treatment was arranged with its own recirculated, UV-sterilized, and dechlorinated water system. During incubation, water changes of approximately 15 gallons were completed each day. Water quality tests for ammonia, nitrates, and nitrites were completed weekly, while pH, dissolved oxygen and temperature were monitored daily using HOBO® temperature loggers (ONSET®) and a LabQuest®2 unit (Vernier Connections™). The daily average temperature from the HOBO® logger was calculated to determine accumulated thermal units (ATU) which allows for the standardization between temperature systems (See Appendix B), as development and growth in fish is highly temperature dependent (Chezik et al., 2014).

### Survival tracking methods

Dead eggs or embryos were cleared from the incubation cells every other day and removed and recorded. Mortalities were sorted into three categories: eyed, hatch, and fry. Stages were chosen based upon relevance to hatchery rearing methods (where eggs are removed if they fail to eye or hatch) and to account for sensitive developmental stages (Hoitsy, 2012). Any unfertilized eggs or those that died immediately post fertilization (as indicated by failure to eye) were considered to be not-eyed. All unfertilized eggs were removed three days post fertilization in the first removal. ‘Eyed’ mortality were fish that died either before hatch or those that never survived hatching. To be considered dead at the hatched stage a fish had to be at least 70% (estimated visually) out of their egg with the shell not restricting their movement. After a minimum of 326 ATU (when the majority of all eggs were eyed based on visual inspection and standardized between temperature treatments) any non-eyed eggs were considered dead and removed. Survival percentage was calculated using the total number of fertilized eggs per replicate (mean: 403 ± 97, Range: 175-688 eggs). The number of eggs per sample was counted from photos of incubation cups using ImageJ multipoint tool (Rasband,1997-2018).

Visible malformations were tracked as a secondary measure of survival as malformed individuals are culled from stocking populations (Branson, 2008; T. W. K. Fraser et al., 2015; Ihssen, 1978). Malformations were categorized as spinal (curved or curled spine), gametal fusion (dicephalous, tricephalous, anadidymus type, katadidymus type, anakatadidymus type, and parasitic), or blue sac disease (Ihssen, 1978; Laale, 1984; Toften & Jobling, 1996; T. S. Yamamoto et al., 1996); with each category representing malformations with both genetic and environmental origins. We also tracked individuals who had more than one malformation type, labelled as “multiple malformations” (**Figure 2)**.

**Figure 1:**
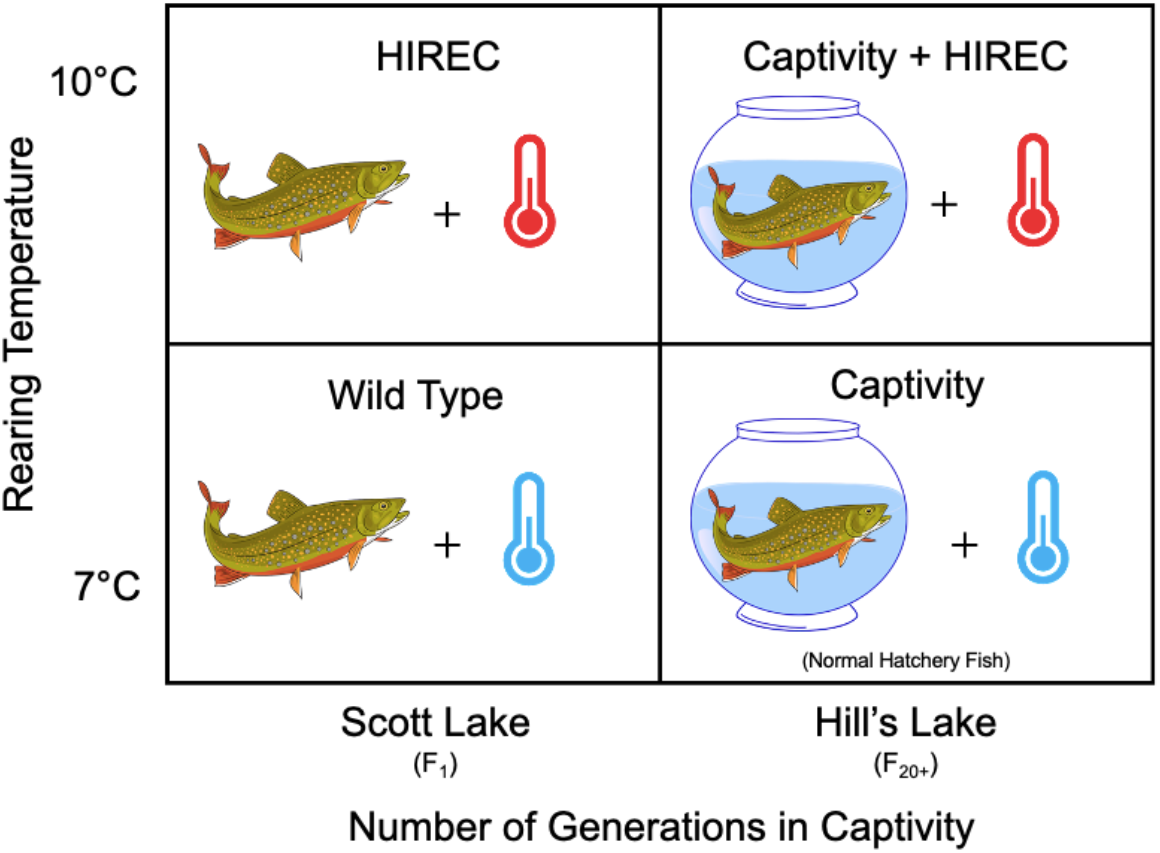
Experimental design to investigate impacts of captivity. Eggs from each family were raised between two temperatures. The experiment was repeated for both populations.

**Figure 2:**
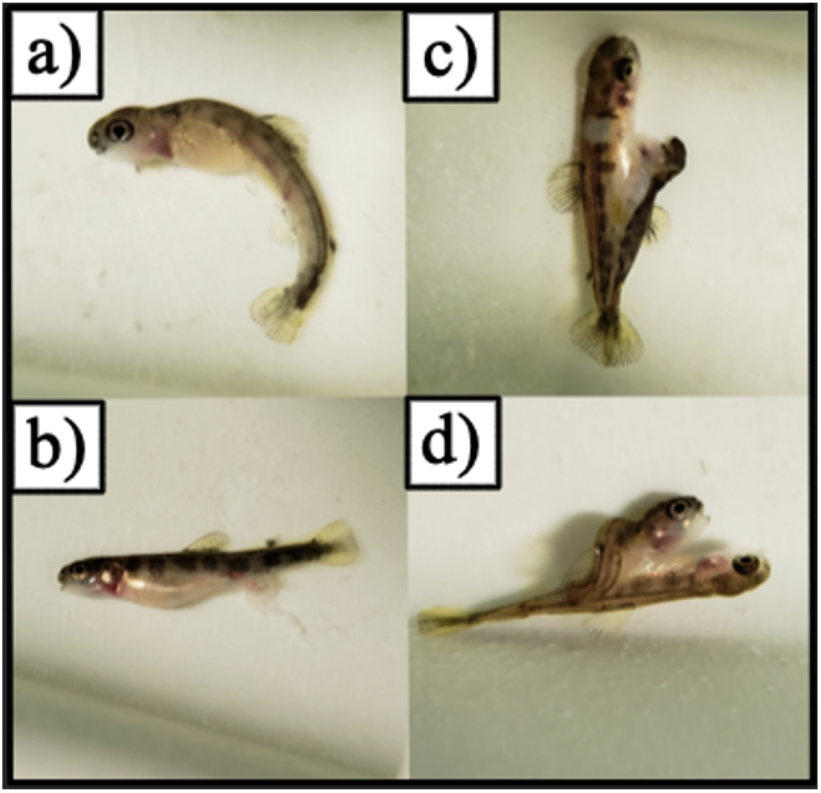
Brook trout try with morphological malformations in the four assigned categories: a) spinal malformations, b) blue sac disease, c) gametal fusion, d) multiple malformations (e.g., spinal malformation and gametal fusion pictured here).

### Statistical Methods

We completed all statistical analyses using R version 4.1.0 (R Core Team, 2021). Using the *full fact* package (Houde & Pitcher, 2016) we performed a binary expansion on survival data at each developmental stage, beginning at fertilized (with non-fertilized eggs removed from analysis), giving individuals alive in a given stage a score of 1 and dead individuals a score of 0. We analyzed this binary survival data with a generalized linear mixed model using a binomial distribution with a log link function, with temperature (categorical: 7°C and 10°C), strain (categorical: F_20+_Hill’s (domestic) and F_1_ Scott (F_1_ Captive)), stage (categorical, 3 levels: eyed, hatch, exogenous feed), strain by temperature interaction, temperature by stage interaction, and strain by stage interaction as main effects. We also included estimated initial egg number per family as a covariate in the model (continuous, range: 341 – 627) to control for density effects. Family identification number (categorical, 1-40 families), female (dam) identity (categorical, 1-20 dams), male (sire) identity (categorical, 1-20 sires), dam ID x sire ID (family) interaction, fertilization block ID (categorical, 1-10 fertilization blocks) and incubation cell identity (i.e., replicate; categorical, 1-12 cells) nested in incubation tray (categorical, 1-14 trays) were included as random effects.

For assessing the likelihood of hatched individuals with developmental malformations we performed the same binary expansion giving “no-malformation” fish a score of 1 and “malformation” fish a score of 0. We analyzed malformation data with a generalized linear mixed model using a binomial distribution with a log link function with temperature, strain, and their interaction as fixed effects, including egg number as a covariate and using the same random-effects structure as above. To analyze the occurrence of specific types of malformations among fish expressing malformations, we used a negative binomial model as the count data were overdispersed. In addition to the fixed- and random effects structure used above, malformation type (categorical: spinal malformations, bluesac disease, gametal fusion, presence of multiple malformations) was included, along with the following interactions: strain by temperature interaction, strain by malformation type interaction, and temperature by malformation-type interaction.

Following the methods defined in Capelle et al. (2017) and Warriner et al. (2020), we analysed model fit using the restricted maximum likelihood approach (Zuur et al. 2009). When a significant interaction was present (p ≤ .05), all interactions were retained and pairwise comparisons between significantly interacting variables were made using Tukey’s HSD. Both significant and insignificant interactions were verified by LRT model comparison using ANOVA. Non-parental random effects were individually removed from the models if they were non-significant, as assessed by the same method. Dam, sire, and family effects were retained regardless of their significance to account for their contributions to overall model variance. We assessed the normality of assumption of model deviance residuals of our binomial regressions by visual inspection of quantile-quantile residual plots and histogram of residuals. Our negative binomial model residuals were additionally assessed using the Kolmogorov-Smirnov test (Hartig 2021). Packages used in performing statistical analyses include: *Fullfact* (Houde and Pitcher, 2021) to convert continuous into binary data, *MASS* (Venable and Ripley, 2002) for liner mixed models and nonlinear least squares, *glmmTMB* (Brooks et al., 2017) to perform negative binomial regression, *lme4* (Bates et al., 2015) and *lmerTest* (Kuznetsova, Brockhoff, and Christensen, 2017) to run GLMMs, *emmeans* (Lenth, 2021) for posthoc comparisons, *DHARMa* (Hartig, 2021) to test for overdispersion and assess non-Pearson residuals, *ggeffects* (Lüdecke, 2018) for model predicted plots, and *ggplot2* (Wickham 2016) for visualizations.

## Results

### Survival Analysis

Number of hatchery generations (strain) significantly interacted with developmental stage to affect survival of Brook trout eggs (p=<0.001 *χ* ^2^=217.8. Temperature and strain had no significant interaction effect on survival (p=0.493 *χ* ^2^= 0.113). Strain and stage significantly interacted to reduce survival (*χ* ^2^ = 217.8, p= <0.001); however, this interaction was driven by differences between irrelevant comparisons (e.g., F_1_eyed - F_20+_ hatch). When restricting post-hoc tests between strains to within-stage comparisons (e.g., F_1_eyed - F_20+_ eyed), we found no difference between probability of survival. **(Table 1)**. Temperature and stage individually had significant effects on survival (Temperature p= 0.028, stage p= <0.01), while strain alone had no effect on embryo survival to exogenous feed (p=0.902). Under elevated temperatures, hatch- and fry stages were more sensitive, exhibiting a decrease in survival in comparison to these stages reared under current conditions **(Figure 3)**. Egg density did not significantly impact the likelihood of survival (p=0.796 *χ* ^2^=0.062). Random effects of dam (*χ* ^2^=21.448 p= <0.001), family (*χ* ^2^=12.306 p=<0.001), and incubation cell nested within tray (*χ* ^2^= 731.93 p= <0.001) significantly contributed to variance in survival. Sire was not a significant random effect (*χ* ^2^=0.485 p=0.4819).

**Table 1:** Percent survival including standard error between treatments and across developmental stages. Sample size (*N*) is reflective of total fertilized eggs for a given treatment group. For samples sizes at each stage see Appendix A

| Strain | Temperature Treatment | Eyed | Hatch | Fry | $N$ |
| --- | --- | --- | --- | --- | --- |
| *F<sub>20+</sub> Hill's* | 7°C | 0.887 ± 0.029 | 0.992 ± 0.002 | 0.995 ± 0.004 | 18239 |
|  | 10°C | 0.865 ± 0.034 | 0.980 ± 0.006 | 0.989 ± 0.003 | 18130 |
| *F<sub>1</sub> Scott* | 7°C | 0.892 ± 0.030 | 0.985 ± 0.006 | 0.988 ± 0.004 | 14934 |
|  | 10°C | 0.866 ± 0.036 | 0.961 ± 0.002 | 0.982 ± 0.006 | 14218 |

**Figure 3:**
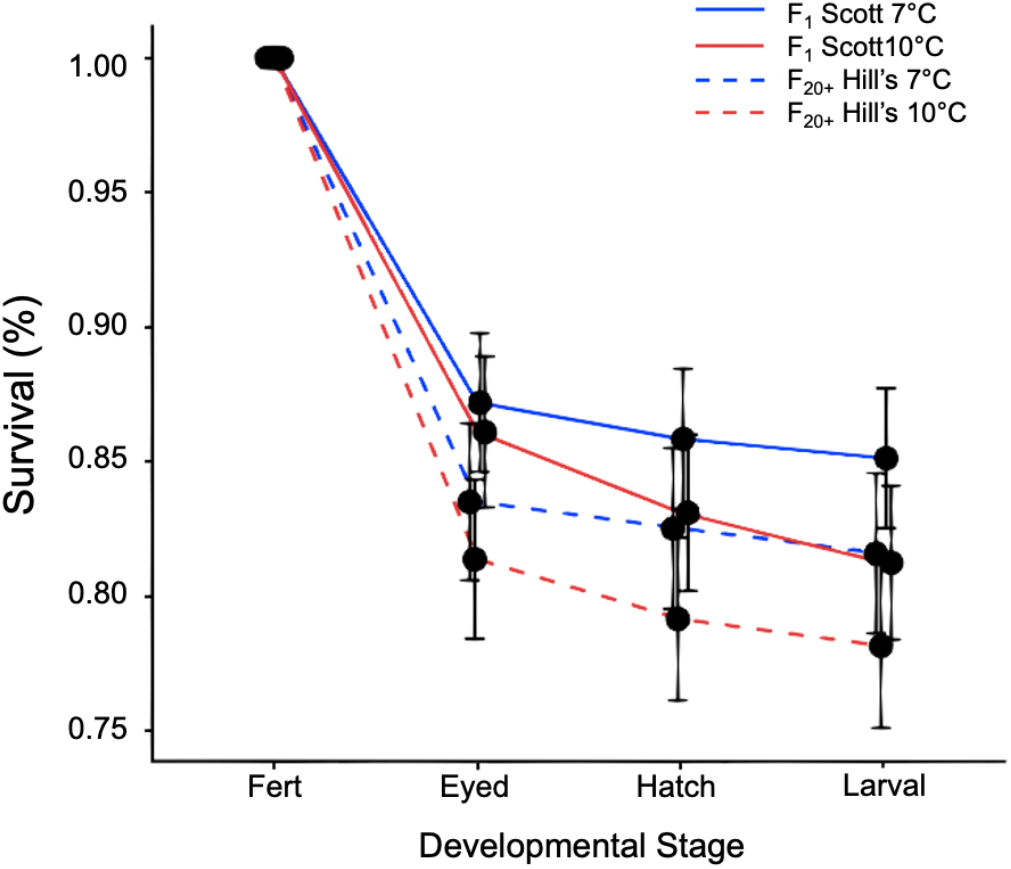
Percent survival across all developmental stages, error bars indicating SE. Elevated temperatures significantly lowered survival between the hatch and fry stages, the eyed stage was unaffected.

### Instances of fry with developmental malformation

Temperature and strain interacted to significantly affect the likelihood of malformations in Brook trout (*χ* ^2^= 5.589 p=0.013). Temperature (p= 0.181) and strain (p=0.272) were not significant as fixed effects **(Figure 4)**. Female egg density had no significant impact on the likelihood of malformations (*χ* ^2^ = 0.626 p=0.419). Random effects of dam (*χ* ^2^ =9.914 p=0.007) and incubation cell nested in tray (*χ* ^2^=149.6 p=<0.001) significantly contributed to variance. Family (*χ* ^2^ = 0.756 p=0.685) and sire (*χ* ^2^ = 1.197 p=0.550). did not significantly contribute to variation Under current temperatures, F_20+_ Hill’s and F_1_ Scott did not experience different malformation likelihoods (Tukey’s HSD: p=0.690). Similarly, each strain experienced similar likelihoods of malformations under elevated rearing temperature (Tukey’s HSD: p=0.913). While F_20+_ Hill’s fish did not experience significant change malformation rates *between* temperature treatments (Tukey’s HSD: p= 0.539), F_1_ Scott fish did experience a significant increase in their likelihood of malformations from current to elevated temperatures (Tukey’s HSD: p=<0.001).

**Figure 4:**
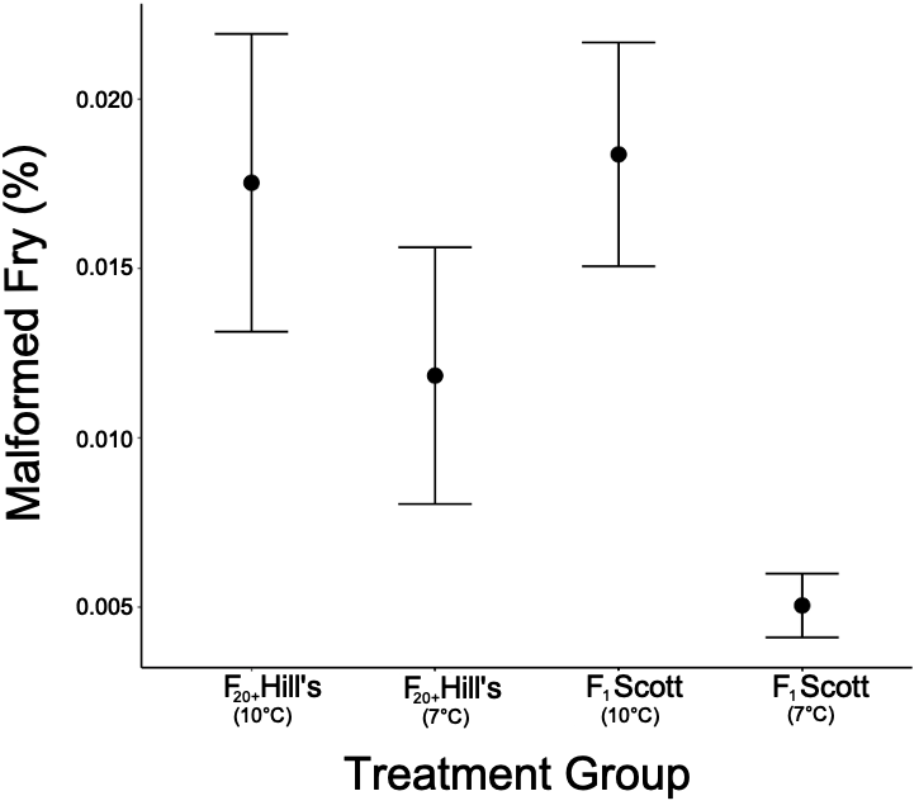
Number of malformed individuals (regardless of type) between treatment groups. F_1_ Scott fish were significantly more likely to exhibit malformations due to increased temperatures.

### Malformations by type

Temperature and strain significantly interacted to influence the number of malformations exhibited (Nb: *χ* ^2^= 8.062 p=<0.001). Type of malformation significantly interacted both with temperature (Nb: *χ* ^2^= 7.846, p=0.020) as well as strain (Nb: *χ* ^2^=20.61 p=0.004) to influence malformations expressed. Temperature regime (p=0.143) and strain (p=0.802) as main effects significantly impacted the type of malformations exhibited **(Figure 5)**. Egg density had no effect on number of malformations (*χ* ^2^=0.066 p=0.797). Replicates (i.e., incubation cell ID nested in tray ID) significantly contributed to model variance (*χ* ^2^=4.366 p=0.037). Random effects of dam (*χ* ^2^=2.379 p=0.123), sire (*χ* ^2^= 0 p= 0.999), and family (*χ* ^2^= 2.275 p=0.132) did not contribute significantly to variance. With respect to the total number of malformations expressed, under both temperature regimes F_20+_Hill’s and F_1_Scott fish showed no significant differences between number of fish with malformations (Tukey’s HSD: F_1_Scott-F_20+_Hill’s 7°C p = 0.291; F_1_Scott-F_20+_Hill’s 10°C p=0.921). However, temperature and strain interactions were significant, with F_1_Scott fish having greater numbers of malformed individuals on elevated temperatures compared to current (Tukey’s HSD: p = <0.001), while F_20+_Hill’s fish showed no significant changes to the number of malformed individuals between temperatures (Tukey’s HSD: p = 0.838). For ease of interpretation of the interaction effects of mutation type, we report further findings within each temperature treatment.

**Figure 5:**
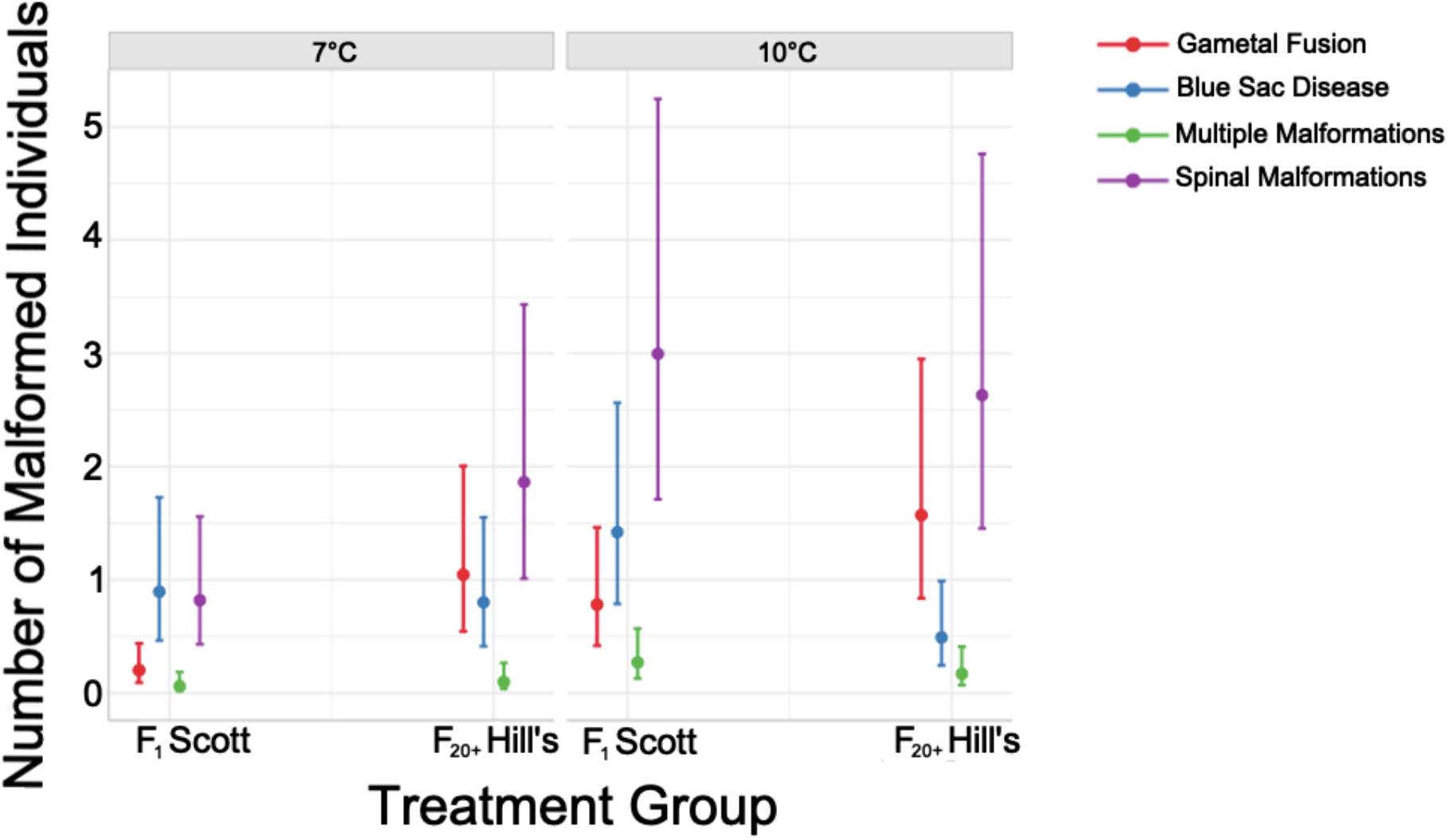
Number of malformed individuals by type for two different strains—F_20+_Hill’s and F_1_Scott, reared on two temperature regimes: elevated (10°C) and Current (7°C). Elevated temperatures increased instances of malformation and the F_20+_Hil’s strain showed greater differential expression of the types of malformations. F_1_Scott fish experienced a greater increase in the number of malformations under elevated versus current rearing temperatures.

### Current Temperature

When reared on current temperatures, F_20+_ Hill’s fish showed no significant differences between specific types of malformations (Tukey’s HSD: blue sac - fusion p=1.00, blue sac - spinal p=0.287, fusion - spinal p=0.843); however, significantly fewer individuals possessed multiple malformations (Tukey’s HSD: blue sac - multiple p=0.004, fusion - multiple p= 0.002, multiple - spinal p=<0.001). F_1_ Scott fish had similar numbers of spinal malformations and blue sac disease (Tukey’s HSD: p= 1.0), both of which occurred more than gametal fusion (Tukey’s HSD: spinal-fusion, p= 0.013 blue sac- fusion p= 0.013) and multiple malformations (Tukey’s HSD: spinal – multiple p=<0.050, blue sac – multiple p= <0.001); and the number of fish exhibiting gametal fusion and multiple malformations did not differ (Tukey’s HSD: p= 2.076).

### Elevated temperature

F_20+_ Hill’s fish reared on elevated temperatures now showed significant differences between malformation types with significantly more individuals having spinal and fusion malformations than blue sac disease (Tukey’s HSD: blue sac - fusion p=0.043, blue sac - spinal p= <0.001); and significantly fewer individuals had multiple malformations than fusion- and spinal malformations (Tukey’s HSD: fusion - multiple p= <0.001, spinal - multiple p=<0.001). There was no significant difference between individuals with blue sac disease and those with multiple malformations (Tukey’s HSD: p=0.593). Similarly, there was no difference between the number of individuals exhibiting spinal malformations and those showing gametal fusion (Tukey’s HSD: p=0.985). For fish from the F_1_ Scott strain raised on elevated temperatures, spinal malformations were more common than gametal fusion and multiple malformations (Tukey’s HSD: spinal-fusion p= <0.001, spinal-multiple p=<0.001) but again no different than blue sac disease (Tukey’s HSD: spinal-blue sac p= 0.260). Blue sac disease was significantly more common than both gametal fusion and multiple malformations (Tukey’s HSD: blue sac-multiple p= <0.001, blue sac-fusion p=0.832), while gametal fusion was not significantly more common an any other malformation type (Tukey’s HSD: fusion-multiple p= 0.240). **(Table 2)**

**Table 2:** Number of malformations by category (including standard error) between treatments and across strains. Sample size (*N*) is reflective of total malformed fry for a given treatment group. For sample of each type see Appendix A.

|  |  | Spinal Malformations | Blue sac disease | Gametal fusion | Multiple Malformations | *N* |
| --- | --- | --- | --- | --- | --- | --- |
| Strain | *F*<sub>20+</sub> Hill's | 2.203 ± 0.592 | 0.624 ± 0.188 | 1.276 ± 0.363 | 0.129 ± 0.051 | 456 |
|  | *F*<sub>1</sub> Scott | 1.56 ± 0.423 | 1.122 ± 0.312 | 0.395 ± 0.125 | 0.128 ± 0.051 | 313 |
| Temperature Treatment | 7°C | 1.230 ± 0.291 | 0.843 ± 0.212 | 0.457 ± 0.128 | 0.077 ± 0.035 | 260 |
|  | 10°C | 2.794 ± 0.593 | 0.831 ± 0.203 | 1.103 ± 0.258 | 0.214 ± 0.07 | 509 |

## Discussion

Climate change is causing increased rates of declines in cold-water adapted species such as salmonids (Myers et al., 2017). To combat the trends in decreasing populations, captive breeding programs have been put into place (Desforges et al., 2022; Rytwinski et al., 2021), yet introduce a new suite of domestication conditions (Farquharson et al., 2021) that either selects for individuals to become optimally adapted to these benign conditions – and subsequently unable to cope with extraneous stressors, or reduces reproductive fitness through various means. Temperature stressors (intentional hatchery practices for conservation and/or production efficiency) along with captive-selection pressures are known to interact and further reduce fitness of salmonids when released into wild settings (Kostow, 2004). In our study, we sought to investigate the interactive effects of rearing temperature and the number of generations in captivity on the survival and instances (and diversity) of malformations in fry brook trout originating from a provincial fish culture station bred for the purpose of reintroductions. The lack of significant difference in egg survival between strains suggest fewer generations in captivity may not improve upon the impacts of captivity in the absence or presence of a stressor. In addition, the newly-captive strain exhibited reduced quality of fry fish under elevated rearing temperatures. With these findings, we posit that in our study, the benefits of fewer generations in captivity are reduced (D. J. Fraser et al., 2019), and that the domesticated strain, at least at the early-life stage, is more robust to temperature stressors.

### Survival under elevated rearing temperatures

In accordance with our original predictions, elevated rearing temperature significantly reduced the likelihood of survival, and its effects were seen predominantly between the hatched and exogenous feed (fry) stages. Overall, we found a decrease in survival from 83.5% at current 7°C incubation temperatures to 79.7% at elevated 10°C temperatures, an overall decrease of 3.8%. Previous Brook trout studies have similarly found decreased rates of survival with elevated rearing temperatures (temperatures used range from 9.4-24°C; Baird et al., 2002; Hokanson et al., 1973; Marten, 1992); in some instances, maximum survival rates have ranged from 77-90% with an average maxima of 83% on optimal (current) rearing temperatures (Hokanson et al., 1973). Given that early life and development are universally thermally sensitive (Dahlke et al., 2020), it is possible that our lack of strain/captivity time effect is a consequence of Brook trout experiencing too great of a thermal bottle neck at these temperatures to be able to detect differences in strain survival.

Elevated rearing temperatures above a species thermal optimum are known to reduce survival in Brook trout (Baird et al., 2002; Hokanson et al., 1973). Despite this reduction, increased rearing temperatures are often used to increase speed of development and manipulate the timing of fish production (D. J. Fraser et al., 2019; Marten, 1992), induce triploidy and sterilize fish for release (Galbreath & Samples, 2000), and to prepare individuals for post release environments with elevated water temperatures (Cook, Wilson, et al., 2018; Durtsche et al., 2021; Penney et al., 2021; Warriner et al., 2020). It is thought that elevated temperatures drive eggs to hatch in an underdeveloped state, due to accelerated activity and development of the hatch gland, compared to fish reared on temperatures that are cooler and closer to optimal (Hayes et al., 1953); and could be one cause of increased mortality, particularly at the hatched stage as found in our study. Increased metabolic activity is thought to be another factor in the reduction of survival as embryos experience an increased oxygen demand and production of waste products (Cingi et al., 2010). Increased metabolic wastes have also been linked to occurrences of blue sac disease (Madison et al., 2020), and could be another source of mortality for fish reared on elevated temperatures. Nonetheless, while warmer waters have been found to have lower levels of dissolved oxygen (Irby et al., 2018), this variable was controlled for in our study (maintained above 9mg L^-1^ across both treatments (Cook, Burness, et al., 2018)); and water changes occurred every day to minimize waste accumulation. Instead we posit that increased instances of blue sac disease for fish reared on elevated temperatures in our system was reflective of temperature conditions as opposed to another environmental factor. While fertilization success is impacted by elevated temperatures (Cingi et al., 2010), it is important to note that differences in survival in our study were not impacted by temperature at fertilization, as both treatment groups were fertilized with cool and consistent water temperatures before being moved into incubation stacks. Additional studies have found the effects of increased temperatures on survival lessened after fish reach the eyed stage suggesting that manipulating development through rearing temperature after this point should not be harmful for embryos (Hokanson et al., 1973; Marten, 1992); however, our results do not support this assertion as we found significant interaction between temperature and stage with reductions in survival occurring at the hatched and fry stages at elevated temperatures.

### Domestication and egg-to-fry survival

Contrary to our predictions, F_20+_ Hill’s fish did not exhibit higher levels of survival under optimal rearing conditions compared to F_1_ Scott, lending no support to captivity optimization in our strains. Furthermore, contrary to our predictions, F_1_ Scott fish did not fare better on elevated rearing temperature (indicative of increased plasticity from more recent generational exposure to unpredictable environments). We found a significant interaction between developmental stage and strain; however, controlling for temperature effects, significance was driven by irrelevant comparisons between stages (i.e. 7°C F_1_Scott eyed – 10°C F_20+_Hill’s fry), thus no meaningful significant interaction was present upon post hoc analysis.

Previous studies have shown that wild fish have greater thermal tolerance than hatchery fish possibly due to their increased genetic diversity and retaining plasticity due to their naturally varying environment (Carline & Machung, 2001). Our results do not support these finding for our system s as we found no interactions between strain and temperature on survival, suggesting that the F_1_ Scott strain was as equally as sensitive as the F_20+_ Hill’s strain under elevated temperatures (as well as current temperatures), and may not possess the requisite plasticity (in terms of thermal tolerance) when faced with this stressor. Recent research has shown that one generation in captivity is able to elicit phenotypic change resulting in no phenotypic differences between F_1_ and F_>1_ generations of captive fish (Farquharson, 2020; Gossieaux et al., 2020). Therefore, our results partially align with the findings of Fraser et al. (2019), suggesting domestication (and unintentional selection for traits unsuitable to the wild) can possibly occur within one captive generation. Previous reviews have also identified phenotypic changes in one generation (Mason et al., 2013) but, as outlined in Milla et al. (2021), true domestication requires multiple generations for domestic characteristics (e.g. growth and nutritional needs, reproductive performance, immune function, and stress response) to emerge fully; essentially, rates of phenotypic change due to domestication are trait specific. Based on lack of strain-temperature interactions and strain fixed effects, it is possible that early offspring survival as well as thermal tolerance are some of the first phenotypic changes brought about by captivity.

### Temperature and strain interact to increase the instance and types of malformation

Previous studies of salmonids (e.g., Crichigno et al., 2021; McDermid et al., 2010) have found malformation rates of over 1% to 14%; our fish showed lower rates of malformation with a maximum rate of approximately 0.02%. We found no significant difference in the likelihood of malformations between strains of Brook trout under either temperature regime, which was contrary to our predictions. Past studies have found hatchery populations to have higher levels of malformations in more domesticated individuals (e.g., Lake trout (McDermid et al., 2010)); however this was not the case with our fish.

Instead, elevated rearing temperatures differentially impacted the F_1_ Scott strain, with the newly captive generation being more sensitive to elevated-than current temperatures, having a greater likelihood of malformed individuals and total number as well. Our results do not support our predictions that F_1_ Scott fish would be more robust and exhibit greater plasticity when faced with environmental stressors. It also does not support our predictions that F_20+_ Hill’s fish would be of lower quality overall due to either the accumulation of genetic deleterious alleles or from a loss of plasticity from prolonged times in captivity as they did not exhibit a loss of quality or increased thermal sensitivity.

Elevated temperatures coupled with genetic factors are known to bring about various types of malformations (Branson, 2008); and hatchery managers experience loss of biomass not necessarily in numbers but in quality when considering individuals with developmental malformations (Ihssen, 1978). While it is not recommended that families producing high numbers of malformed individuals be integrated into broodstock (Sneddon et al., 2016), hatchery managers may not be afforded the luxury of removing breeding adults as it has been shown that low numbers of breeding adults in captive populations can reduce both genetic and phenotypic diversity which may have negative effects on the wild populations that are being supplemented (O’Sullivan et al., 2020; Perez-Enriquez et al., 1999).

Our study was one of the first to investigate the prevalence of various types of malformations in salmonids as current literature tends to focus on one specific type of malformation or malformations in one area of the body (i.e., vertebral malformations, head and jaw malformations, or twinning). In our study, spinal (or vertebral) malformations were generally more common than other categories across temperature and generation time; these malformations have been found to have a strong genetic basis as well as being caused by nutritional deficiencies (Branson, 2008). Spinal malformations have varying degrees of severity and often do not impact growth and survival, but may impact condition factor and severely impact quality of produced fish (T. W. K. Fraser et al., 2015). With no reduction in growth and survival, it is possible families producing large numbers of fish with spinal malformations go unnoticed, contributing to the persistence of the responsible genotypes within domesticated breeding stocks as egg quality is often determined by egg survival (Bromage et al., 1992; Olk et al., 2019). Additionally, spinal malformations showed significant increases under elevated rearing temperatures, irrespective of strain, suggesting a strong environmental driver of this malformation type.

Instances of twinning (and rare but still-present triplets), which we categorized as gametal fusion, were more common in our highly captive strain under both temperature regimes. Similar to spinal malformations, twinning is thought to also have a strong genetic basis and these greater proportions could have been a result of unintentional inbreeding within the hatchery system (Nowosad & Kucharczyk, 2019). Nevertheless, elevated temperatures have been used to induce twinning in fish (e.g. Atlantic salmon (Fjelldal et al., 2016)), and studies consistently show a higher instance of fusion malformations in heat-treated groups (Owusu-Frimpong & Hargreaves, 2000; T. Yamamoto et al., 1998). Higher instances of this gametal fusion in highly domestic fishes suggest the need to investigate existing brood stocks still displaying these developmental malformations and determine if genetic, environmental, or GxE interactions are the most likely source (Christie et al., 2016). Finally, blue sac disease is a common phenomenon that occurs when fish experience a metabolic change in response to environmental conditions like elevated temperatures (Madison et al., 2020). Although its expression was not influenced by temperatures in our study, it is a known occurrence in Brook trout that has been found to have bacterial (*Aeromonas hydrophila*) and environmental causes such as crowding stress and nitrogenous wastes (Kayiş et al., 2015).

Our findings of F_1_ Scott fish exhibiting both a higher likelihood of malformation and greater number of malformations when reared under elevated temperatures suggest that Scott fish are more thermally sensitive than the longer-captive strain in terms of fry development. It is thought that increased handling stress on eggs, as well as elevated temperatures, can disrupt genetically determined developmental patterns and morphology (Brown et al., 2010). The F_1_ generation may therefore be currently experiencing a fitness reduction due to acclimation to captive settings; in essence, they have yet to adapt to increased disturbance of hatchery care and rearing densities (Kayiş et al., 2015). Interestingly, F_20+_ Hill’s strain did not show an inability to cope with a novel environmental stressor, indicated by lower thermal sensitivity in fry development. A possible explanation is the origins of the Hill’s Lake strain occupied historical latitudes lower than Scott Lake fish, and therefore could be considered a warm-adapted strain (see Cook et al., 2018b). However, the recent Hill’s Lake strain shares mixed genetic material with the Scott Lake strain (Stitt et al., 2014), and exhibited no difference in performance with respect to survival regardless of thermal regime. Additionally, Hill’s Lake fish are typically used in studies as the representative “domesticated” strain in comparison to others (e.g., Howell et al., 2022; McDermid et al., 2012; Stitt et al., 2014). Taking these factors into consideration, we believe our results to be mostly explained by F1 Scott strain’s lack of acclimation capacity. Nevertheless, future studies should consider investigating thermal sensitivity of an intermediately-captive (F1< F_n_ < F_20+_) strain from the same source population to determine if this thermal sensitivity decreases with time and adaptation occurs to other confounding hatchery stressors.

What is particularly interesting about our results is the differential expression of malformation types between strains. F_1_Scott fish showed significantly more larvae with blue sac disease than rates of gametal fusion, as well as significantly more larvae with spinal malformations than gametal fusion, while F_20+_Hill’s fish showed no differences at all between these same types. Additionally, F_1_Scott fish showed no significant difference in the number of larvae with blue sac disease and spinal malformations while F_20+_Hill’s fish had significantly more individuals with spinal malformations than blue sac disease. It is possible that these differences are driven by the increased thermal sensitivity that has been shown by F_1_Scott fish. While our study only considered malformations that were easily identified upon visual inspection, future research should consider detailed imaging of fry to determine if malformations not easily identified visually are present (i.e. cardiovascular, head and jaw, and gill malformations (Branson, 2008)). A more complete analysis of malformed individuals to identify possible genetic sources of congenital malformations within brood stocks would allow for the breeding of higher quality individuals.

### Maternal effects in egg survival and quality

We found significant maternal effects on the variation in survival and instances of malformations in our Brook trout strains. A study by Penney et al. (2018) investigating embryonic phenotypic plasticity in Brook trout found maternal effects to outweigh abiotic factors (e.g., pH and fluctuating water temperatures) with respect to egg quality and survival. Maternal effects have previously been found to also have significant impacts on salmonid progeny survival and quality (as defined by body size in this example), whereas sire effects or family interactions were found to be more population- and stage specific (Houde, Black, et al., 2015). The same study by Houde et al (2015a) also determined maternal environment to be one of the important factors pertaining to early life fitness and survival. Comprehensive evaluation of dam performance and reproductive output in breeding programs is often difficult and costly to do (Bobe, 2015); however, maternal identity significantly contributing to our model variance indicates the potential for identifying negative effects of broodstock dams on egg survival. As common measures of maternal quality such as the relationship between egg size and offspring survival do not hold true for Brook trout - due to possible habitat constraints making the energetic costs of large eggs outweigh the survival benefits to large offspring (Baficinar, 2004; Hutchings, 1991), future studies should replicate the work of those such as Khendek et al. (2017) and investigate the physiological impacts of captivity on oogenesis and reproductive performance in captive Brook trout to determine the mechanistic reasons for strong maternal effects in progeny survival. Because fish in early life stages are entirely dependent upon the environment and resources supplied by their parents, including genetic makeup and egg quality (e.g. lipid content, egg size) (Burt et al., 2011), this large investment from dams to their eggs is a probable reason for finding significant maternal effects but no significant sire or consistent family effects (the latter being indicative of non-additive effects). Our findings along with others highlight the importance of investigating drivers of survival at early life stages as understanding these relationships can allow conservation managers to adapt practices to exploit biological processes and assist cold-adapted species in adapting for future environments (Cook, Wilson, et al., 2018; Peck et al., 2012; Warriner, 2019).

## Conclusion

With concerns for the effects stocked fishes have on the production in natural systems (Einum & Fleming, 2001; Ersbak & Haase, 1983; Gossieaux et al., 2020; Kostow, 2004), characterizing the implications of warming water temperatures on the survival and quality of stocked fish both in hatchery settings, the wild, and with varying generations spent in captivity (as our study addressed), is imperative for predicting the success of such programs (Einum & Fleming, 2001; O’Sullivan et al., 2020; Venditti et al., 2018). As with other studies (Acornley, 1999; Beacham & Murray, 1985; Bellinger et al., 2018; Chezik et al., 2014; Cingi et al., 2010; Marten, 1992; Molony et al., 2004; T. S. Yamamoto et al., 1996) our own has shown that elevated temperatures affect offspring survival of cold-adapted freshwater salmonids. However, we additionally demonstrate certain developmental stages to be more sensitive to warmer temperatures, significantly reducing survival at the hatch and fry stages. We also show this stressor to equally affect early survival of different strains of contrasting generation times, but to have a greater impact on fry quality of newer captive strains. Breeding fish with varying generational captivity times has been suggested as a mitigation effect for domestication selection (Milla et al., 2021). Our results imply that this practice may not provide the assumed fitness benefits of greater captivity- or thermal stressor capacity in newly captive strains. Furthermore, the use of elevated temperatures to increase growth rates, prime offspring, and create multiple stocking events should be utilized with caution as it may come with non-insignificant production-efficiency and conservation costs.

## Supporting information

Supplemental Appendicies

## Acknowledgements

The authors would like to thank the staff at the Codrington Fish Culture Station, University of Windsor animal care staff, Semeniuk Lab members, Dr. Oliver Love, Love lab members, GENFish for funding.

