## Supplemental Appendicies for "Effects of generations in captivity and elevated rearing temperature on Ontario hatchery brook trout (*Salvelinus fontinalis*) fry quality and survival"

### APPENDICES

*Appendix A*

| | ***Strain*** | | | |
|---|---|---|---|---|
| | *$F_{20+}$ Hill's* | | *$F_1$ Scott* | |
| ***Temperature Treatment*** | **7°C** | **10°C** | **7°C** | **10°C** |
| Fert | 18239 | 18130 | 14934 | 14218 |
| Eyed | 14949 | 14478 | 12846 | 11988 |
| Hatch | 14760 | 14038 | 12616 | 11503 |
| Fry | 14585 | 13848 | 12487 | 11171 |
| Normal | 14395 | 13580 | 12417 | 10930 |
| Malformed | 188 | 268 | 72 | 241 |
| Gametal Fusion | 81 | 115 | 9 | 50 |
| Blue Sac Disease | 31 | 20 | 29 | 68 |
| Spinal | 84 | 141 | 33 | 139 |
| Multiple | 6 | 8 | 1 | 18 |

**Appendix A.** Sample sizes between treatment groups, for each: developmental stage, malformed individuals, and malformation category. Malformation counts may not be equal to the total number of fry fish, as some fish were able to escape from incubation cups and unable to be counted in survival analysis.

*Appendix B*

| | ***Fertilization Date*** | | | | | | | |
|---|---|---|---|---|---|---|---|---|
| | *Nov. 15 2019* | | *Nov. 22 2019* | | *Nov. 29 2019* | | *Treatment Average* | |
| ***Temperature Treatment*** | **7°C** | **10°C** | **7°C** | **10°C** | **7°C** | **10°C** | **7°C** | **10°C** |
| Eyed | 371.1 | 316.2 | 370.9 | 348.4 | 364.3 | 364.4 | 368.8 | 343.0 |
| Hatch | 520.9 | 499.1 | 514.3 | 521.3 | 520.6 | 506.8 | 518.6 | 509.1 |
| Fry | 981.9 | 946.6 | 955.7 | 897.6 | 922.8 | 842.5 | 953.5 | 895.6 |

**Appendix B.** Accumulated thermal units at which fish were considered eyed, hatched, and fry between treatment groups.
